# Visual cues for moisture perception of facial skin: Enhancing high-frequency components of skin lightness increases perceived dryness

**DOI:** 10.1101/2024.07.16.603839

**Authors:** Yuya Hasegawa, Hideki Tamura, Tama Kanematsu, Yuzuka Yamada, Yohei Ishiguro, Shigeki Nakauchi, Tetsuto Minami

## Abstract

Facial skin texture provides crucial visual cues that reflect an individual’s impressions and health conditions. In this study, we focused on the texture attribute of “moisture” and investigated which visual cues influenced skin moisture perception. The stimuli consisted of images from three facial areas (the whole face, cheek, and eyebrow areas) with and without makeup under two lighting directions. The participants rated the presented stimuli on the three texture attributes (moisture, glossiness, and attractiveness) using a five-point scale. The results from Experiment 1 revealed correlations between the ratings and histogram statistics of each channel in the CIELAB color space, with variations depending on the conditions and facial regions. A negative correlation was observed between the cheek moisture perception and the variance in the *L\** channel. We subsequently obtained similar ratings by enhancing the high-frequency components of skin lightness for artificially dried stimuli (Experiment 2) and for images depicting different skin conditions due to various types of makeup (Experiment 3). Both experiments confirmed a decrease in moisture and attractiveness and an increase in glossiness; these were correlated with the degree of artificial drying. These findings indicated that the high-frequency components of skin lightness could be visual cues for determining the perceived dryness.

## 1. Introduction

Facial skin provides essential visual information that reflects a person’s impression and health condition. For example, the relationship between skin condition and age is well documented [1– 5] and linked to health [6] and attractiveness [7]. In addition to long-term changes in age and health, short-term changes in blood flow also affect facial impressions. For example, the skin appearance physiologically changes, and blood flow and oxyhemoglobin serve as cues that indicate emotions and health status [8,9]. Thus, increased skin redness can strongly express anger [10–14]. In support of this, anger expression influences the memory of the red color [15]. Similar to emotions, increased skin redness also affects “attractiveness” [16,17]. Additionally, for Japanese women’s faces, a reddish complexion is perceived as whiter if the brightness is constant [18]. Subsequent international comparative experiments indicate that ethnicity and the environment influence this effect [19]. These changes in skin color impact the unconscious processing of emotions [20]. Thus, these studies highlight the importance of the skin as a signal to others.

Higher-order information obtained from the skin, such as texture, is also considered to contribute significantly to the overall impression of the skin. Specifically, the effect of skin gloss (radiance) on perception has frequently attracted attention [21,22]. For example, when facial attractiveness is evaluated, people consider the reflection of light from the skin [23]. Additionally, a previous study revealed that attractiveness increased in the order of nonglossy “matte,” negatively perceived glossiness “oily shiny,” and positively perceived glossiness “radiance” when facial attractiveness was judged [24]. Moreover, image statistics that contribute to the perception of skin gloss have been identified, showing that humans use features such as variance and skewness of luminance as cues to judge the glossiness of object surfaces [25,26]; this correlates with the perceived age of the skin [27,28]. Matsubara et al. suggested a relationship among the skin reflection, subsurface scattering, and image statistics [22]. Furthermore, a recent study comprehensively investigated the relationships between the various skin textures and image statistics, beyond only glossiness. Otaka et al. aimed to extract the principal components of skin texture from ratings based on nine adjectives for skin textures. They reported that the skin texture could be explained by a surface interpreted by the two axes of pleasant and glossy, which correlated with image statistics [29]. Based on the results from these studies, the visual system uses certain cues that can be predicted by image statistics to estimate the skin texture.

Here, we focused on the largely unknown aspect of how the visual system perceives “skin moisture” among various skin textures. The physical quantity of skin moisture reflects both the healthiness of moisture regulation and the amount of moisture retention [30,31]. These factors indicate resistance to dryness and pathogens [32]. Additionally, the sensation of a person’s skin changes with the surrounding environmental temperature and humidity [33]. Thus, visually estimating the moisture in another person’s skin is important for assessing health conditions. When the skin is moisturized, the light scattering on the skin surface decreases, allowing more light to penetrate deeper into the skin. Thus, the skin appears darker, pinkish, and translucent [34]. Even though moisture was included as a factor in Otaka et al.’s study mentioned above, it was positioned diagonally away from both the pleasant and glossy axes, unlike other texture adjectives [29]. This finding indicated that skin moisture had a more complex texture than the other texture attributes. Additionally, while studies on the wetness perception on object surfaces [35] and stain perception [36] have been performed, these studies involve actual or simulated wet conditions. In contrast, skin moisture refers to the perceived level of hydration of the skin, even without a visible liquid on the surface or a wet sensation upon touch. Due to these factors, the determination of how the visual system perceives skin moisture is challenging.

Therefore, the aim of this study is to elucidate the visual cues that the visual system uses to estimate the texture of skin moisture. We conducted three psychophysical experiments in which participants rated the texture of images of human skin (including manipulated images) as the experimental stimuli. In Experiment 1, we investigated the differences in the ratings related to image statistics for various skin images with respect to condition, area, and shooting direction. This enabled the identification of the candidate image statistics associated with skin moisture. We found a correlation between the moisture and the variance in lightness. Based on these findings, in Experiment 2, we proposed an image processing method that manipulated the image statistics associated with moisture and investigated the effect of these changes on the moisture ratings. In Experiment 3, we applied this method to a different set of images to confirm the robustness of the cue and examined the relationship between the visual cues and physical quantities of the skin.

## 2. Experiment 1

In Experiment 1, we investigated the visual cues contributing to the perception of skin moisture.

### 2.1 Methods

#### 2.1.1 Observers

Twenty students (10 males and 10 females; average age: 23.00 ± 3.23 years) from Toyohashi University of Technology and Kyushu University participated in Experiment 1. This exploratory experiment did not involve a priori power analysis, and the sample size was determined to meet the minimum requirement of at least 10 participants of each gender. To minimize the effect of differing ethnic backgrounds on skin impressions, all participants were of Asian descent, and the experimental stimuli featured Japanese faces (see Stimuli). The participants had normal or corrected-to-normal vision. The experiment was conducted with the approval of the Ethics Review Committee for Experiments Involving Human Subjects at Toyohashi University of Technology and adhered to the Declaration of Helsinki. Written informed consent was obtained from all participants prior to the experiment.

#### 2.1.2 Apparatus

The experiment occurred in a dark room. Stimuli were displayed on an EIZO CG319X or EIZO EV2450 monitor, with a resolution of 1920 × 1080 pixels and a frequency of 60 Hz. Each monitor was calibrated to a white point of D65 and a gamma of 2.2.

#### 2.1.3 Stimuli

Images of the skin of 23 Japanese women (13 in their 20s, 1 in their 40s, 3 in their 50s, and 6 in their 60s) were used as experimental stimuli (Fig. 1A); these were collected by a camera (NIKON D850) in a room with blocked external light and a white cloth background. Illumination was provided from the front using a super lamp holder SLH3 (Andoer), and the faces were photographed from a frontal angle of 0 degrees or an oblique angle of 45 degrees. Images were collected under two conditions: with and without makeup. Afterward, they were cropped to exclude hair, ears, and necks, with a focus on the entire face (Fig. 1B). Further cropping created square images of the cheek and rectangular images of the eyebrow area. The entire face and cheek are commonly studied in skin texture studies [e.g., 29]. We additionally included the eyebrow area based on the assumption that dryness was more likely to occur in this region. In total, 276 images were used (23 individuals × 2 orientations × 2 makeup conditions × 3 regions). The stimuli were presented within a visual angle of 14.25° square.

**Fig. 1.**
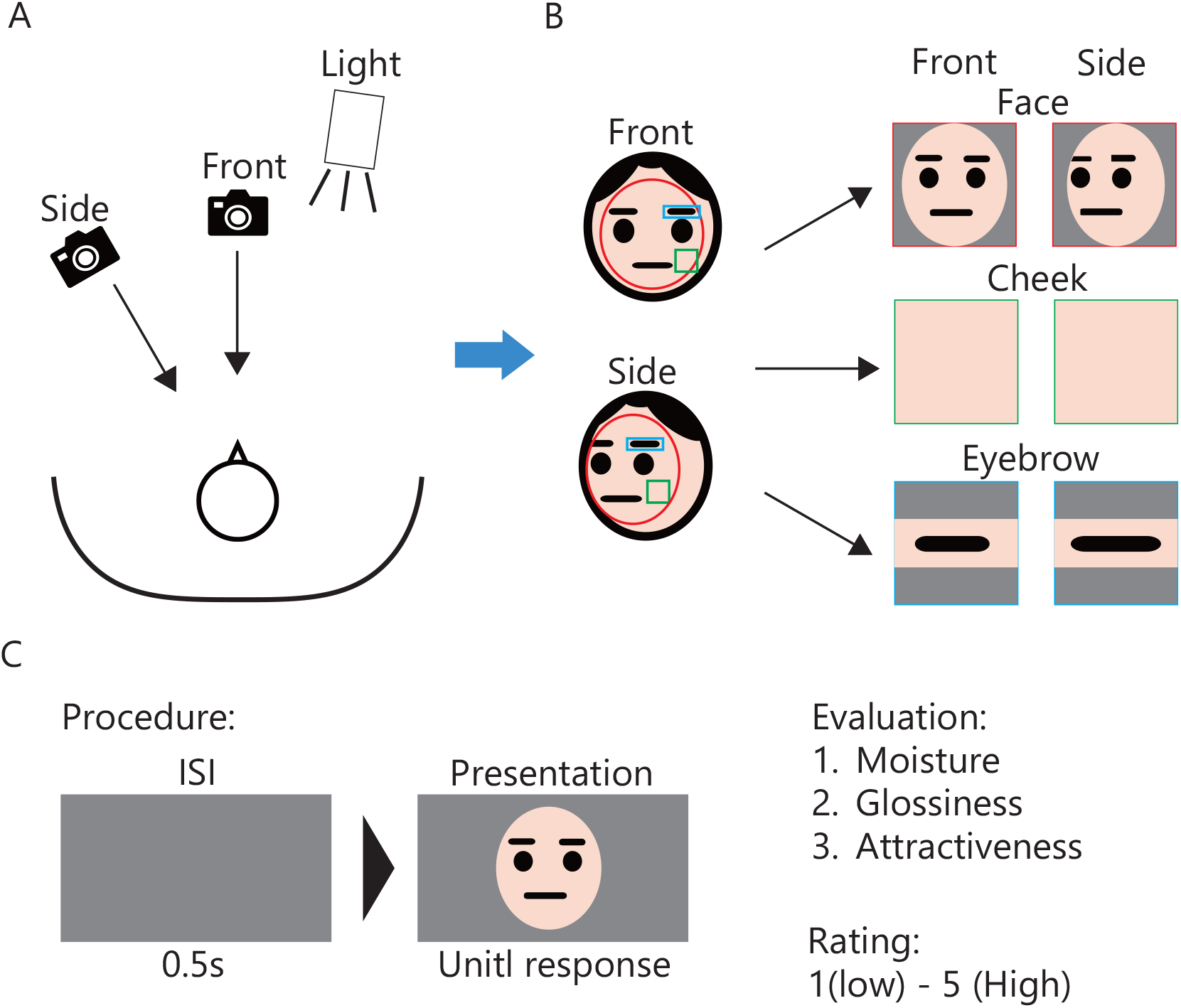
Experimental stimuli and procedure. (A) Photographing conditions. (B) Direction and region conditions of the stimuli. (C) Procedure.

#### 2.1.4 Procedure and tasks

The participants sat 70 cm from the monitor with a chin rest and viewed the stimuli directly from the front. The flow of one trial was as follows (see also Fig. 1C). Each trial began with a 0.5-second interstimulus interval (ISI) with a gray background (*x* = 0.314, *y* = 0.328, *Y* = 20.93), followed by the presentation of a stimulus image, which remained on the screen until a response. The participants were asked to rate the texture of each image for moisture, glossiness, and attractiveness on a 5-point scale, where 5 indicated a high and 1 indicated a low perception of the texture. Moisture was included to achieve the objective of this study, whereas glossiness and attractiveness were included for comparison and validation since they have been commonly used in previous studies.

Each participant completed 828 trials (276 stimuli × 3 textures). The trials were organized into nine blocks, combining the three texture adjectives and three regions (92 trials per block), with the block order counterbalanced among participants. The trial order within each block was randomized. The participants took breaks every 46 trials. The experiment was conducted using MATLAB R2021a and Psychtoolbox 3.0 [37–39].

#### 2.1.5 Data analysis

Data analysis was performed via MATLAB R2024a. The ratings from the participants were averaged for each condition. Initially, the ratings for moisture, glossiness, and attractiveness were calculated by averaging across shooting directions and makeup conditions for the facial, cheek, and eyebrow regions of the 23 individual facial stimuli (Fig. 2).

**Fig. 2.**
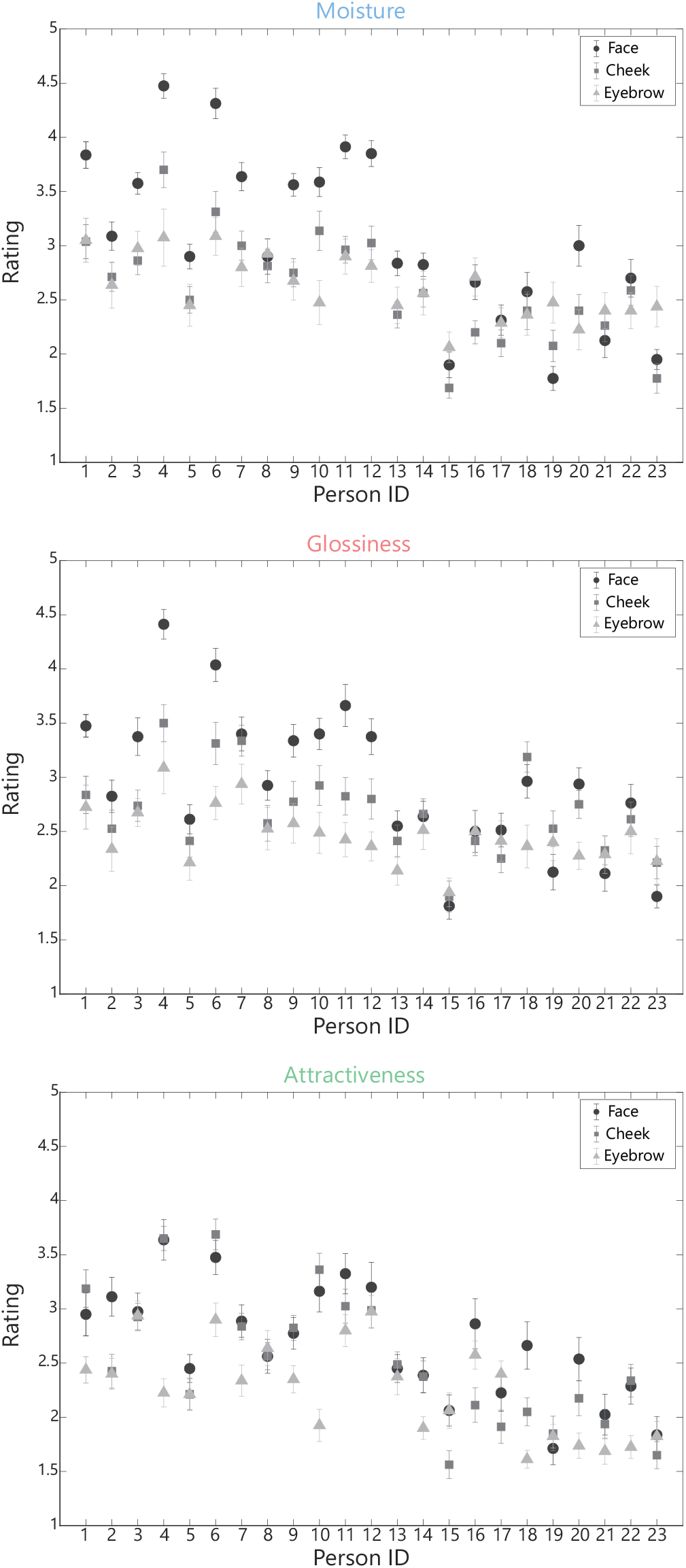
Ratings. Each panel shows different attributes (glossiness, moisture, and attractiveness). The horizontal axis indicates the ID of individuals in the generation order from 20 s to 60 s. The vertical axis indicates the rating score. The error bars represent the standard errors among the observers.

Next, we analyzed the effects of shooting direction and region and the effects of makeup presence and region on each texture by two-way repeated-measures analyses of variance (ANOVAs) (Figs. 3A and 3B). The p values for the post hoc tests were adjusted via the Tukey method.

**Fig. 3.**
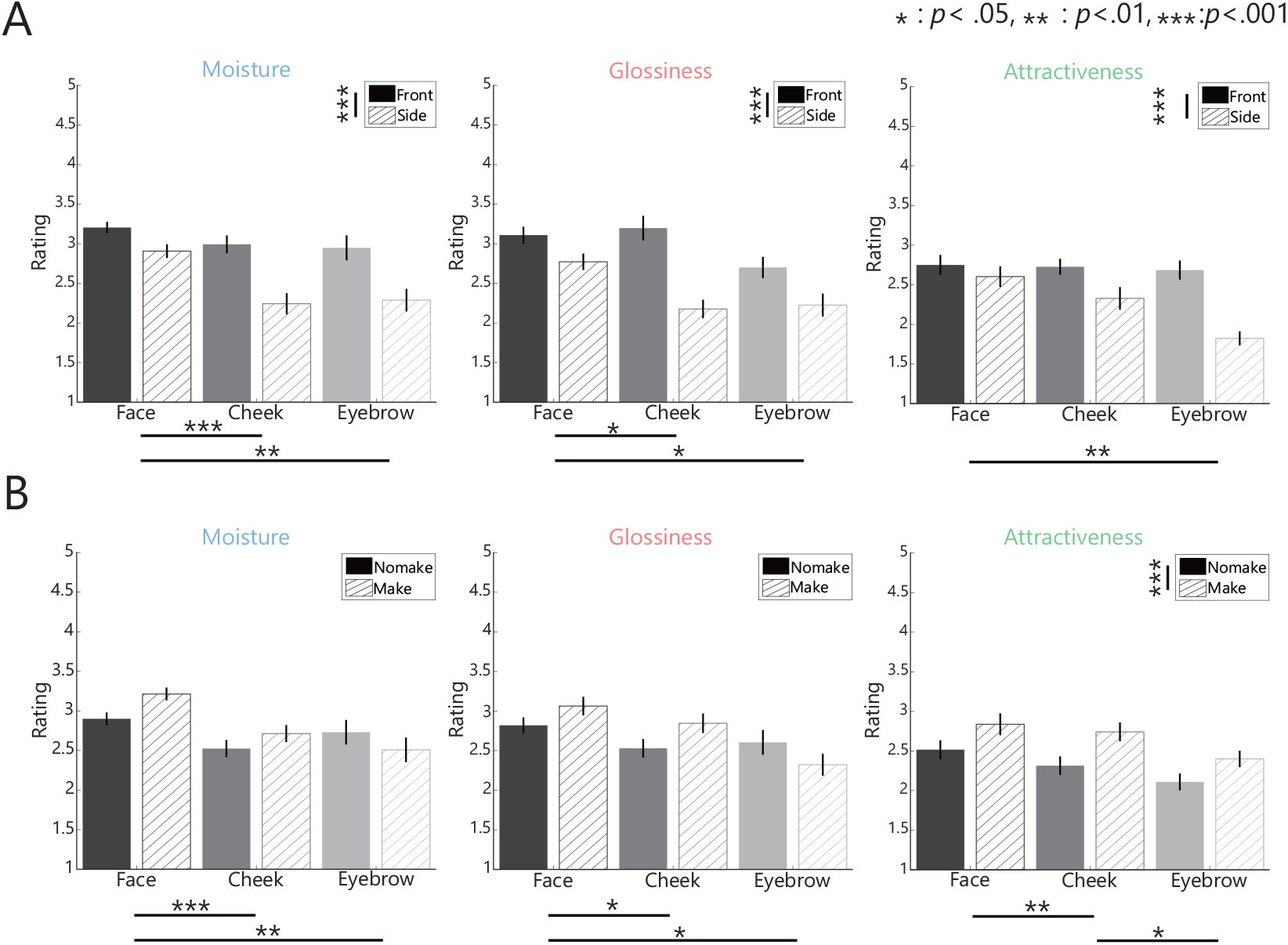
Ratings in terms of direction (A) and makeup (B) The format was the same as that in Fig. 2, except that the horizontal axis indicates the regions and direction (A) or makeup (B) conditions. Asterisks indicate significant differences; *: *p* < 0.05, **: *p* < 0.01, ***: *p* < 0.001.

### 2.2 Results and discussion

#### 2.2.1 Ratings

Fig. 2 shows that the perceived moisture level decreased with increasing age; this result was in agreement with the decrease in stratum corneum lipids with age [40], which may contribute to dryness. In contrast, while a decrease in physical skin moisture (increased transepidermal water loss) with age was observed, this aspect was found to have only a small correlation with visual moisture [41], indicating that the visual perception of moisture may not necessarily correspond directly to the physical quantity.

Similarly, a trend toward decreasing ratings with increasing age for glossiness and attractiveness was observed, and this result was supported by previous findings [2,27]. In the comparisons between the regions, the faces had higher ratings for all texture attributes than the cheek and eyebrow regions. These results indicated that the visual features needed to rate texture attributes were more easily captured when the entire face was viewed.

Next, we focused on moisture and examined the effects of shooting direction on the ratings (Fig. 3A; complete results in Tables S1-12). The direction factor had a significant effect, with “Front” having significantly higher ratings than “Side” (*F*(1,19) = 105.04, *p* < .001, *η*_*p*_^2^ = .847). Thus, the participants perceived the texture more strongly when it was observed from the front. Similarly, a significant main effect of the region factor (*F*(1.28, 24.33) = 9.61, *p* = .003, *η*_*p*_^2^ = .336) was observed; moreover, across texture attributes, “Face” was rated higher than “Eyebrow” (*t*(19) = 3.30, *adj. p* = .001, Cohen ′s *d* = 0.74). Additionally, an interaction effect was observed for all texture ratings (*F*(1.76,33.52) = 8.75, *p* < .001, *η*_*p*_^2^= .315), indicating that in terms of glossiness and moisture, “Face” was rated higher than “Cheek” (*t*(19) = 7.56, *adj. p* < .001, Cohen ′s *d* = 1.69). These results indicate that when the texture of the face and cheeks were evaluated, the face could be perceived with stronger texture sensations since other regions of the face could be referenced.

Similarly, we verified the effects of makeup presence on the ratings (Fig. 3B; complete results in Table S13-21). A significant main effect of the makeup factor was observed only for attractiveness, with significantly higher ratings for “Make” than for “Nomake” (*F*(1,19) = 93.21, *p* < .001, *η*_*p*_^2^= .831). This finding confirmed that attractiveness could be enhanced by altering the appearance of the skin through makeup. Additionally, a significant main effect was observed between the regions, with “Face” being rated higher overall than the other regions (*F*(2.00,37.94) = 6.81, *p* = .003, *η*_*p*_^2^ = 0.264). However, no significant difference was observed between “Face” and “Eyebrow” in terms of attractiveness.

In the “Eyebrow” region, the ratings for moisture and glossiness were higher in the “Nomake” condition (moisture: 2.73 ± 0.70 (Nomake) vs. 2.51 ± 0.70 (Make), glossiness: 2.60 ± 0.70 (Nomake) vs. 2.32 ± 0.62(Make)). This result could be attributed to the eyebrows in the “Eyebrow” condition; this could create a significant visual contrast with the skin, potentially influencing these ratings. However, since the observation of eyebrows alone is rare and the visual impression they create is poorly understood, this aspect remains speculative.

#### 2.2.2 Relationships between the ratings and color statistics

The determination of the visual cues that are used to assess the perception of moisture needs to be examined. To investigate this, we calculated the correlation coefficients between the ratings and the color statistics proposed in previous studies [42]. Similarly, we converted the sRGB values of each image to the CIELAB color space using D65 as the white point (MATLAB “*rgb2lab*” function). Then, we computed the first-to third-order histogram statistics for *L*, a\**, and *b\**: mean, variance, and skewness. Furthermore, we calculated the correlation coefficients between *L\**-*a*, L\**-*b\**, and *a\**-*b\**, resulting in 12 statistics (moisture in Fig. 4; glossiness and attractiveness in Fig. S1).

**Fig. 4.**
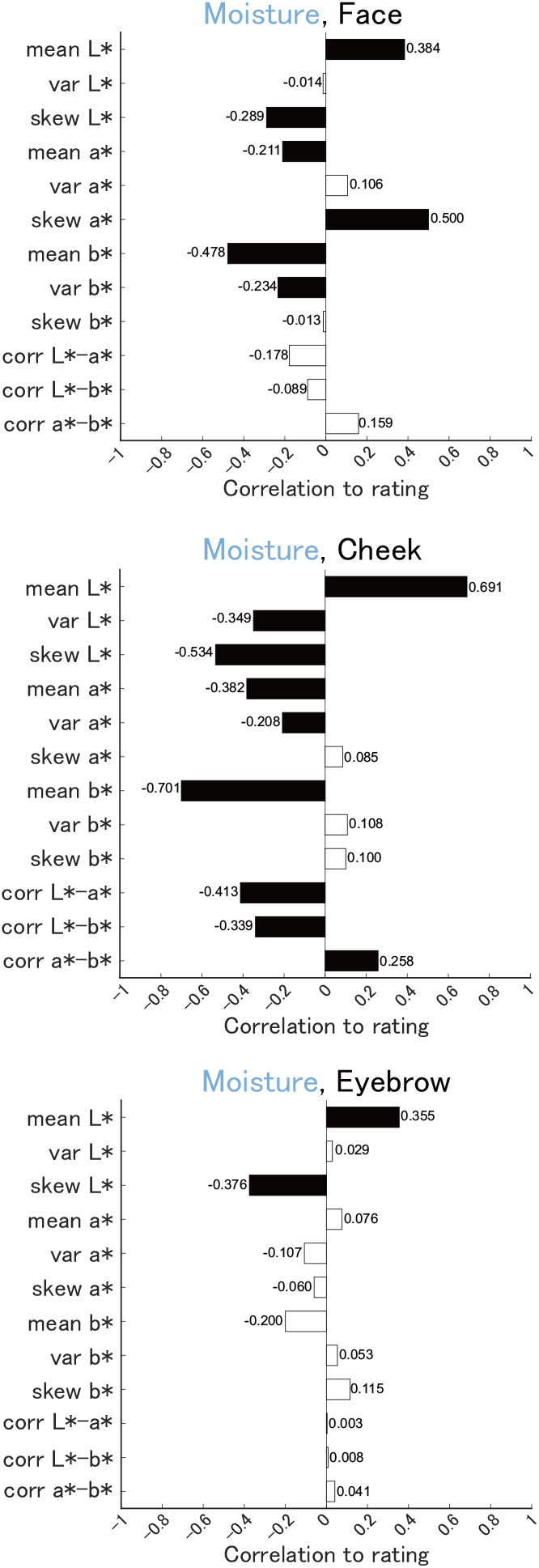
Correlations between the moisture ratings and color statistics in each region (whole face, cheek, and eyebrows).

Fig. 4 shows the correlations between the moisture ratings of the cheeks and the color statistics for each region. We found significant correlations regardless of region, with the mean *L\** showing a positive correlation and the skewness *L\** showing a negative correlation. These results indicated that brighter images indicated by a higher mean *L\** had a higher moisture content; thus, the change in diffuse reflection on the skin surface was associated with hydration. Furthermore, a higher skewness of lightness was associated with lower moisture levels (dryness). This indicated that as the skin became drier, the distribution of light became more skewed.

Focusing on the effect of the skin alone, we found a significant negative correlation in the variance *L\** for the cheek; this result potentially indicated that the lightness distribution widened when the skin was dry. Considering the contribution to pure moisture perception when no regions other than the skin were present, this color statistic was a key candidate cue. Additionally, negative correlations with moisture evaluations were confirmed for the mean *a\** and mean *b\** in the cheeks. These results indicated that greater redness and yellowness were related to lower moisture, which was consistent with the fact that redness increased with dryness [43]. Moreover, the significance of the three statistics of correlations were likely associated with a significant correlation of the mean *L*, a\**, and *b\**.

Thus, based on these findings, the visual system could use a combination of lightness and color distribution cues to estimate the skin moisture. In particular, the variance in lightness appeared to be a primary indicator of the skin area, whereas the mean and skewness of lightness provided additional information. The observed correlations aligned with known physiological changes in skin appearance due to hydration levels, indicating that these visual cues could effectively inform moisture perception.

## 3. Experiment 2

Based on the results of Experiment 1, the perception of skin moisture could be linked to color statistics with respect to the changes in lightness. If this is true, then images processed to alter lightness should induce perceptual changes in moisture sensation, and correspondingly, the color statistics associated with moisture cues should similarly change. Therefore, in Experiment 2, we tested whether the sensation of skin moisture changed by manipulating the lightness appearance through image processing of cheek images.

### 3.1 Methods

#### 3.1.1 Observers

Twenty-one students from Toyohashi University of Technology and Kyushu University (10 males, 11 females; mean age 22.71 ± 3.32 years) participated. The sample size exceeded the criteria used in Experiment 1. The recruitment criteria, ethical review, and participant consent procedures were the same as those in Experiment 1.

#### 3.1.2 Apparatus

The apparatus was the same as that in Experiment 1.

#### 3.1.3 Stimuli

We used cheek images from a frontal view without makeup in Experiment 1 for image processing. Specifically, these images were processed to emphasize the appearance of dryness by enhancing the visibility of white streaks on the skin; these are known signs of dehydration that appear when moisture levels are insufficient [43–45]. The presence or absence of these white streaks influences changes in the lightness of image features, as indicated in Experiment

1. Therefore, we performed image processing to enhance the high-spatial-frequency components of the lightness in the images to emphasize the skin streaks (see Fig. 5A). The steps are as follows.
2. The CIE *L\** values for each pixel in the image are obtained.
3. A 2D frequency Fourier transform (FFT) is performed on the image, ranging from 0 to 2 × pi.
4. The frequency domain of step 2 is adjusted to range from ^−^pi to pi (centering the DC).
5. A frequency domain mesh is created for the image FFT.
6. The distance, *r*, is determined from the DC. The distance from the center to each edge is 1, and the distance to each corner is sqrt(2).
7. Only the frequency components of pixels where the *r* value is above a set threshold are extracted (original = 1, low = 0.2, middle = 0.1, high = 0.01). This method uses a high-pass filter to allow high-frequency components around the periphery.
8. The frequency domain is restored from step 6 to range from 0 to 2 × pi.
9. The spatial domain is converted back using the 2D IFFT.
10. The pixels in step 8 are replaced with values below the mean with the mean value.
11. The values from step 9 (containing only high-frequency/high luminance components) are added to the *L\** values from step 1.
12. The *L\** values from step 10 are applied to generate the final processed image.

**Fig. 5.**
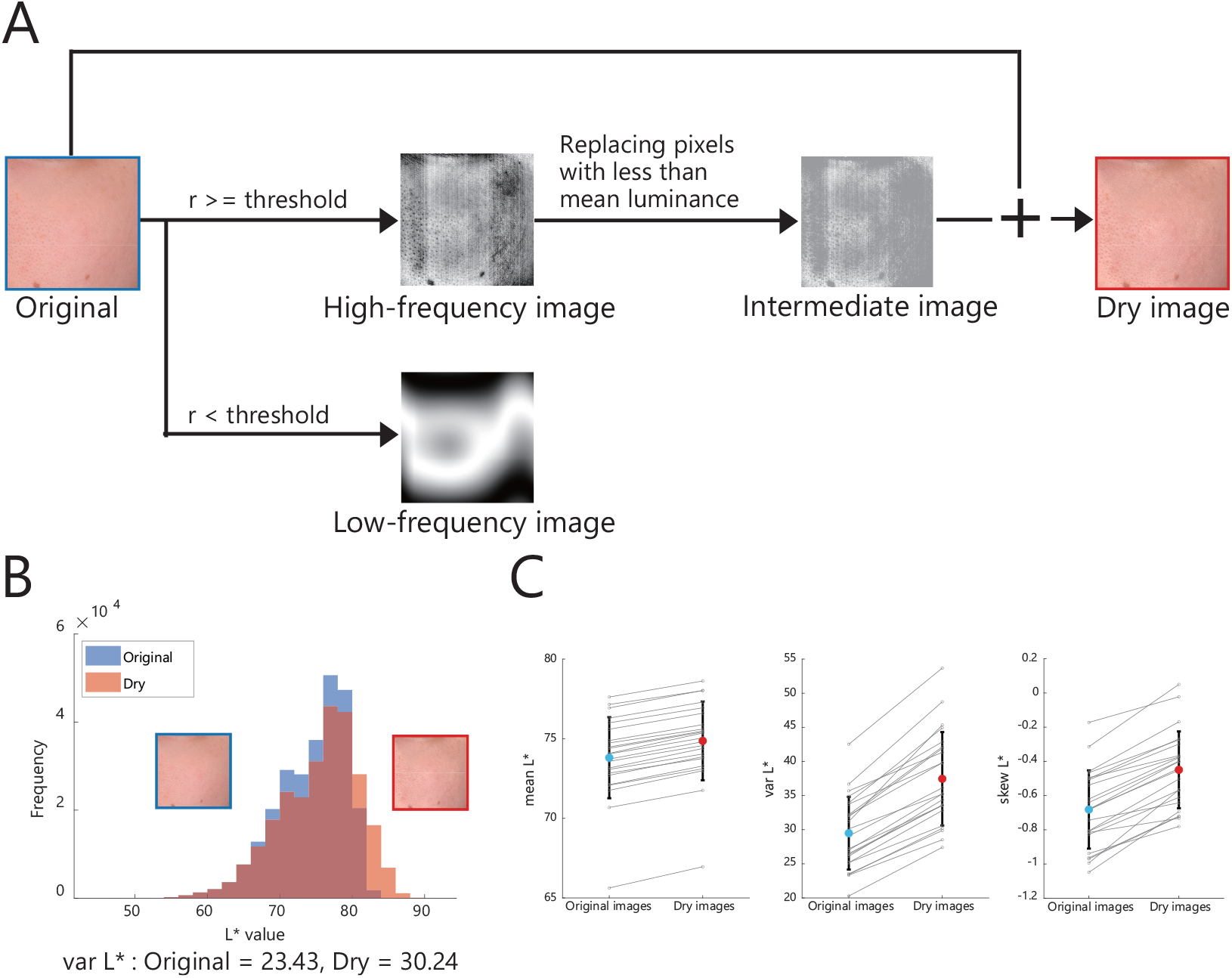
(A) Flowchart of the simulated drying process. (B) Changes in the *L\** histogram of an image after drying. The original var *L\** increased from 23.43 to 30.24. (C) Changes in the *L\** statistics of 23 cheek images before and after drying. From left to right: Mean *L\**, var *L\**, and skew *L\**. Var *L\** and skew *L\** increased significantly due to processing; these results are consistent with the negative correlation between perceived moisture in Experiment 1 and these statistics, whereas the mean *L*\* also increased.

After the pseudo-drying process, we confirmed that the histogram statistics of *L\** shifted to the right (Fig. 5B). We observed an increase in the variance of *L\** in all 23 samples; this increase was associated primarily with the skin dryness (Fig. 5C center). Similarly, the skewness of *L\** also increased (Fig. 5C right), whereas the mean *L\** showed an increasing trend (Fig. 5C left).

#### 3.1.4 Procedure and tasks

The experimental procedure and task were the same as those in Experiment 1. Each participant completed 276 trials, consisting of 23 cheek images × 4 dryness conditions (original, low, middle, and high) × 3 textures. The three texture ratings were divided into separate blocks, and the order of block implementation was counterbalanced across participants. The order of the trials within each block was randomized. The participants took breaks every 92 trials.

#### 3.1.5 Data analysis

The mean ratings were calculated for each image processing condition. We performed two-way repeated-measures ANOVA for the dryness and rating attribute factors.

### 3.2 Results and discussion

Fig. 6 shows that the perceived moisture significantly decreased as the degree of image processing increased (*F*(2.36,47.25) = 29.28, *p* < 0.001, *η*_*p*_^2^ = 0.594). These findings indicate that pseudo-drying could reduce the perceived moisture due to image processing. Additionally, attractiveness decreased with increasing moisture (*F*(2.55,51.05) = 74.95, *p* < 0.001, *η*_*p*_^2^ = 0.789). A significant correlation was observed between moisture and attractiveness (*r* = 0.791, *p* < 0.001), showing a strong association between them. Conversely, glossiness significantly increased with increasing dryness (*F*(1.32,26.42) = 24.18, *p* < 0.001, *η*_*p*_^2^ = 0.547). These results indicated that the modified image features, which were mainly brighter pixels with high-frequency components, contributed to the glossiness by being perceived as specular highlights [46–48].

**Fig. 6.**
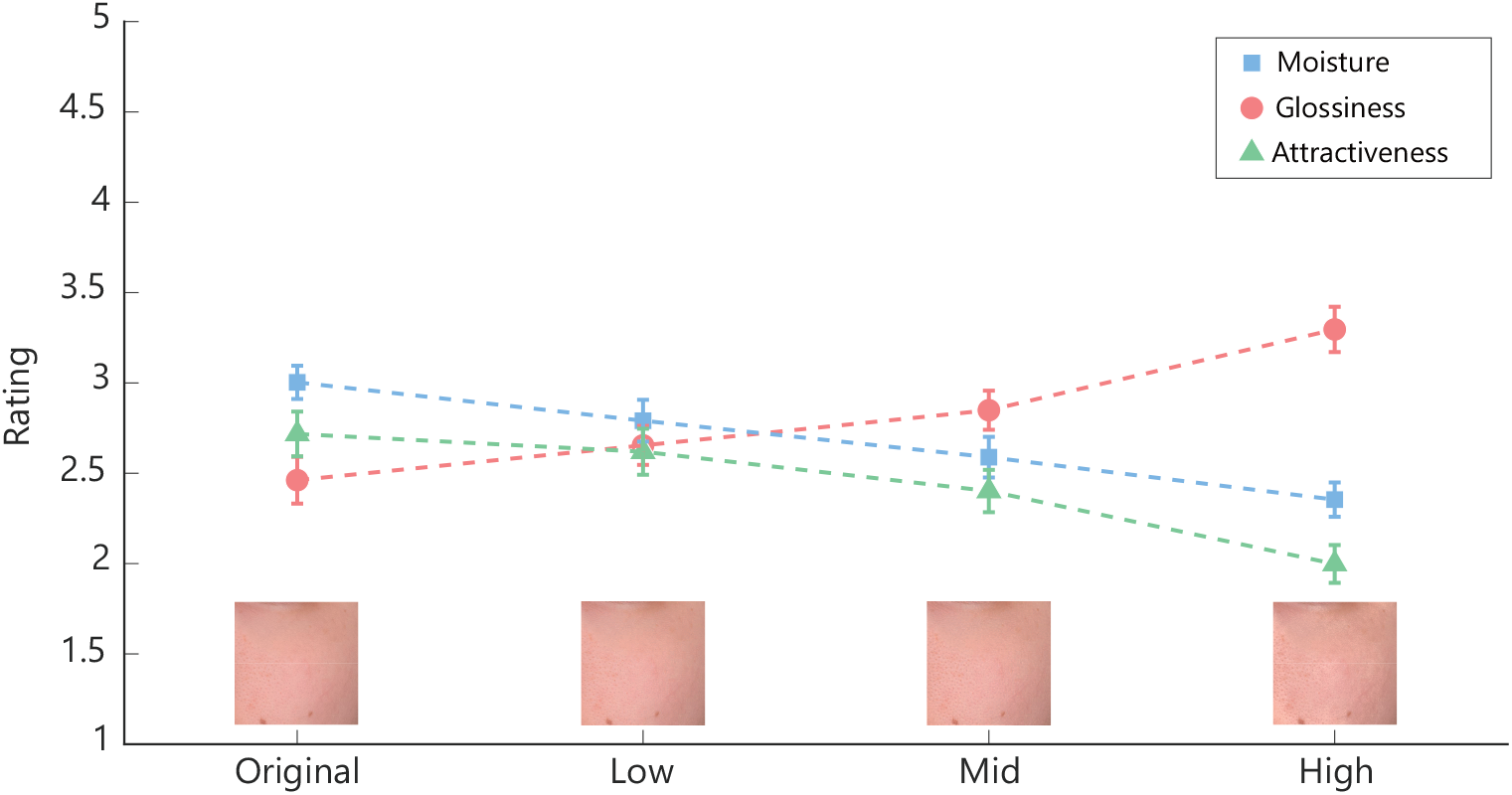
Ratings of Experiment 2. The format was the same as that in Fig. 2, except that the horizontal axis indicates the extent of the image manipulation.

## 4. Experiment 3

Experiment 3 aimed to further verify whether pseudo-drying processing contributed to the perception of dryness by altering image features. If these modified image features determine the perception of moisture and dryness, then new images processed in this manner should also be perceived as drier, regardless of the image set. Therefore, we created a new set of stimuli and applied the same drying process to examine how the ratings changed. Simultaneously, we investigated the relationship between the ratings and physical properties of the skin. We acquired physical measurements of the skin when the new stimuli were captured and analyzed the relationships between the images and the ratings, as well as between the physical properties and the ratings.

### 4.1 Methods

#### 4.1.1 Observers

The experiment involved 20 students from Toyohashi University of Technology and Kyushu University (10 males, 10 females; mean age 22.40 ± 3.05 years), following the same criteria as those used in Experiment 1. All other aspects remained consistent with those of

Experiment 1.

#### 4.1.2 Apparatus

The apparatus was the same as that in Experiments 1 and 2.

#### 4.1.3 Stimuli and skin physical parameters

Two female models in their 20s participated, and the shooting and measurements were conducted every afternoon for four days, approximately once a week (1st, 9th, 15th, and 23rd days), to vary the skin condition due to the biorhythm. After the face was washed each day, four physical quantities were measured: 1) transepidermal water loss (TEWL) by using VAPO SCAN AS-VT100RS (Asahi Techno Lab), 2) stratum corneum water content (SKICON) by using SKICON-200EX-USB (IBS), 3) stratum corneum water content (CORNEO) by using Corneometer CM825 (Courage+Khazaka electronic), and 4) oiliness (OIL) by using Sebumeter SM815 (Courage+Khazaka electronic). The measurement position was the meeting point of a vertical line from the eye’s outer corner and a horizontal line from the nose’s tip. Thus, eight data points were obtained from 2 models × 4 days for each physical quantity (see Table S22). We predicted that the TEWL would negatively correlate with the moisture rating, whereas the SKICON and CORNEO would positively correlate with the moisture rating. Additionally, we expected that the OIL would positively correlate with the glossiness rating.

For the same two models, skin images were collected from the front (0 degrees) and at a 45-degree angle in the same environment as that used in Experiment 1. The images associated with the abovementioned physical quantities were collected at an angle, resulting in 8 images per model after the faces were washed. Additionally, to capture various skin conditions, 12 more images with different skin states were collected from each model’s angle (see Table S23). Therefore, 64 images were collected in total and consisted of 2 models × 2 angles × (4 + 12) types.

Whole-face and cheek images were trimmed from these images using the same method as in Experiment 1. The image processing was then performed on cheek images with the same intensity as the high-intensity images in Experiment 2. A total of 192 images were used in the experiment: 64 whole-face images, 64 cheek images, and 64 dried cheek images.

#### 4.1.4 Procedure and tasks

The rating task was conducted as in Experiments 1 and 2. The number of trials per participant was 576 (192 images × 3 textures). The participants were divided into facial blocks and cheek (original image + dryness image) blocks. The allocation of blocks and rating attributes was counterbalanced across participants. The participants took a break after every 64 trials.

#### 4.1.5 Data analysis

Similar to Experiment 2, we computed the average rating values for each condition, and performed two-way repeated-measures ANOVA. Additionally, the correlations between the physical measurements and ratings were computed.

### 4.2 Results and discussion

As shown in Fig. 7, a significant decrease in moisture content was observed in the cheek images processed with the image manipulation (*F*(1.29, 24.60) = 27.26, *p* < 0.001, *η*_*p*_^2^ = 0.589), and the moisture content of the dried cheek was the lowest (Face-Dry Cheek: *t*(19) = 5.93, *p* < 0.001, Cohen’s *d* = 1.33; Cheek-Dry Cheek: *t*(19) = 5.15, *p* < 0.001, Cohen’s *d* = 1.15). Similarly, we found a significant decrease in the attractiveness (*F*(1.85, 35.22) = 59.87, *p* < 0.001, *η*_*p*_^2^ = 0.759; Face-Dry Cheek: *t*(19) = 9.07, *p* < 0.001, Cohen’s d = 2.03, Chee0k-Dry Cheek: *t*(19) = 11.26, *p* < 0.001, Cohen’s *d* = 2.58) and a significant increase in glossiness (*F*(1.72, 32.74) = 16.01, *p* < 0.001, *η*_*p*_^2^ = 0.457; Face-Dry Cheek: *t*(19) = ™1.17, *p* = 0.484 Cohen’s *d* = ™0.26; Cheek-Dry Cheek: *t*(19) = ™4.97, *p* < 0.001, Cohen’s d = ™1.11); these results were consistent with those from Experiment 2. Therefore, the method proposed for manipulating moisture perception demonstrated a certain degree of robustness across stimulus sets.

**Fig. 7.**
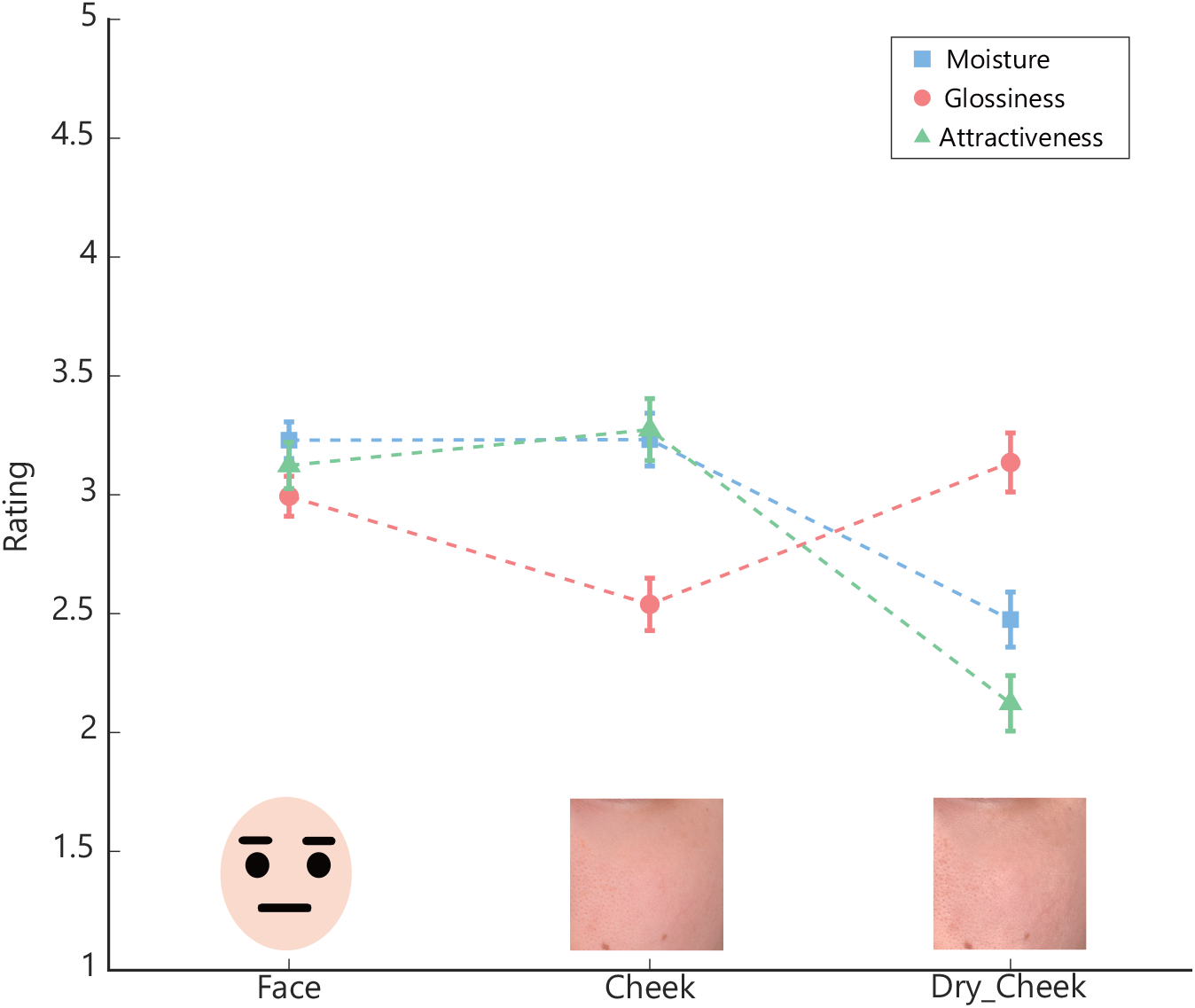
Ratings of Experiment 3. The horizontal axis indicates the image regions or conditions. The other formats were the same as those in Fig. 6.

Table 1 provides the correlation coefficients between the skin ratings and the measured physical quantities (see scatter plots in Fig. S2). As a result, no significant meaningful correlations were observed. Thus, estimating the perceptual texture of the skin solely based on measured physical quantities is challenging, and subjective evaluations are needed to obtain accurate skin impressions.

**Table 1.**
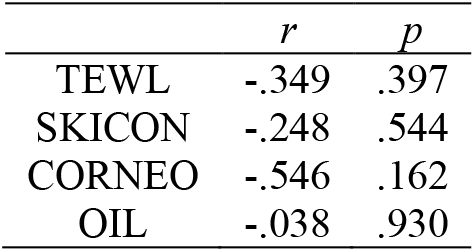
Correlations between the physical quantities and moisture ratings.

## 5. General discussion

The aim of this study was to elucidate the visual cues contributing to the perception of skin moisture. The ratings from Experiment 1 indicated that the sense of moisture decreased with age and that the entire face was more likely to be perceived as moist than just a part of the face. Additionally, correlation analysis between the ratings and color statistics revealed strong correlations between the skin moisture and lightness-related statistical measures. Experiment 2 revealed that images with brighter high-frequency components resulted in reduced moisture perception. Experiment 3 indicated that similar effects appeared when these image features were applied to other sets of skin images and that moisture perception did not necessarily correlate with the physical moisture content of the skin.

These results indicate that modulating the high-frequency components of skin light contributes to the perception of moisture. This aligns with physiological knowledge indicating that white streaks appear on the skin surface due to dryness [43–45]. Additionally, the indirect effect of increasing the local lightness may have affected the perception of skin moisture by increasing contrast, causing the relatively darker areas to appear even darker. This darkening effect also corresponds to physiological dryness events such as the widening of pores or increased visibility of blackheads [49]. Furthermore, wet skin is known to appear darker and exhibit intensified pink coloration [34]. The difference between the results from this study and those from previous studies could be attributed to the difference between being wet and perceiving moisture.

Our findings indicate that changing the lightness of the skin not only contributes to the perception of moisture but also influences other textural attributes. The increase in glossiness is likely attributed to the perception of increased highlights due to lightness. In situations resembling a “truly wet state,” where a liquid is applied to the surface of an object, specular reflection occurs on the liquid surface, resulting in locally brighter regions [35]. This relationship between the skin moisture and glossiness may have contributed to these results.

Since the entire face and the cheeks were presented with the same field of view, the pixels for the entire face were compressed. Consequently, differences in the perception of fine details might have been reflected in the evaluations. Additionally, the results indicate that the recognition of a face as a face [50] may not only be influenced by skin tone [51] and brightness [52,53] but also affect texture perception. Therefore, even simply flipping the face upside down could change the perception of moisture or texture.

We recognize and discuss several limitations of our research. First, with respect to the clues discovered in this study, increasing the high-frequency components of the lightness led to decreased perceived moisture. However, the effect was insufficient when attempting to “increase” the perception of moisture by decreasing this lightness. Thus, some form of asymmetry in the perception of moisture exists, and further research is needed to elucidate the visual cues associated with increasing the perception of moisture.

Second, the results from Experiments 2 and 3 revealed that as the perceived moisture of the skin decreased, the glossiness of the skin increased. This potentially occurred because the increase in lightness reproduced the appearance of white streaks on the skin. Simultaneously, the high-luminance areas were also perceived as specular reflections, leading to an increase in glossiness. Similarly, the simultaneous decrease in perceived attractiveness and moisture showed a negative correlation between attractiveness and glossiness. Therefore, the increase in glossiness observed in this study corresponded to “oily shiny” [23]. To gain deeper insights into the separation of moisture and glossiness, discovering clues that contribute independently to moisture and glossiness is crucial in future research.

Finally, the focus of this study was on the skin of Japanese women, with texture evaluations also conducted by Asian men and women from the same cultural background. Since they belong to the same ethnocultural category, the evaluation scores would not likely show variations due to different ethnicities or cultures. However, in this study, verification targeting male skin was not conducted; thus, our findings need to be verified to determine their generalizability.

## 6. Conclusion

The result from this study revealed that increasing the high-frequency components of skin lightness leads to a decrease in perceived skin moisture; thus, the skin appears drier. Based on the results, these features can serve as cues for perceiving skin dryness.

## Funding

This work was funded by the Japan Society for the Promotion of Science (JSPS) KAKENHI and Pias Corporation.

## Acknowledgments

This work was supported by JSPS KAKENHI (Grant Numbers JP22K17987, JP20H05956, and JP20H04273), and Pias Corporation.

## Disclosures

Y.I. is employed by Pias Corporation. There are no other competing interests to declare.

## Data availability

The data underlying the results presented in this paper are not publicly available at this time but may be obtained from the authors upon reasonable request.

## Notes

### Summary of Updates

A few minor points were revised.

