## Supplement information for "Visual cues for moisture perception of facial skin: Enhancing high-frequency components of skin lightness increases perceived dryness"

### Exp. 1: Front-Side Moisture

Table S1. *Summary of two-way repeated-measures ANOVA of Rating in Front-Side (Moisuture)*

| Effect | $\hat{\eta}_p^2$ | $F$ | $df^{\text{GG}}$ | $df_{\text{res}}^{\text{GG}}$ | $MSE$ | $p$ |
|---|---|---|---|---|---|---|
| Part | .336 | 9.61 | 1.28 | 24.33 | 0.42 | .003 |
| Direction | .847 | 105.04 | 1 | 19 | 0.09 | < .001 |
| Part × Direction | .315 | 8.75 | 1.76 | 33.52 | 0.07 | .001 |

Table S2. *Post hoc comparisons - Part*

| Contrast | $\Delta M$ | 95% $\text{CI}_{\text{Tukey(3)}}$ | $t$ | $df$ | $p_{\text{Tukey(3)}}$ |
|---|---|---|---|---|---|

| Contrast | $\Delta M$ | 95% $\mathrm{CI}_{\mathrm{Tukey}(3)}$ | $t$ | $df$ | $p_{\mathrm{Tukey}(3)}$ |
| --- | --- | --- | --- | --- | --- |
| Face - Cheek | 0.44 | [0.29, 0.59]$^{a}$ | 7.56 | 19 | < .001$^{a}$ |
| Face - Eyebrow | 0.44 | [0.10, 0.77]$^{b}$ | 3.30 | 19 | .010$^{b}$ |
| Cheek - Eyebrow | 0.00 | [-0.35, 0.35]$^{c}$ | 0.00 | 19 | > .999$^{c}$ |

Table S3. *Post hoc comparisons - Direction*

| Contrast | $\Delta M$ | 95% CI | $t$ | $df$ | $p$ |
| --- | --- | --- | --- | --- | --- |
| Front - Side | 0.57 | [0.45, 0.69] | 10.25 | 19 | < .001 |

Table S4. *Post hoc comparisons - Part * Direction*

| Contrast | $\Delta M$ | 95% $\mathrm{CI}_{\mathrm{Tukey}(6)}$ | $t$ | $df$ | $p_{\mathrm{Tukey}(6)}$ |
| --- | --- | --- | --- | --- | --- |
| Face Front - Cheek Front | 0.21 | [-0.01, 0.44] | 2.99 | 19 | .070 |
| Face Front - Eyebrow Front | 0.26 | [-0.17, 0.68] | 1.92 | 19 | .419 |
| Face Front - Face Side | 0.30 | [0.18, 0.42] | 7.84 | 19 | < .001 |
| Face Front - Cheek Side | 0.96 | [0.62, 1.30] | 8.87 | 19 | < .001 |
| Face Front - Eyebrow Side | 0.92 | [0.48, 1.35] | 6.67 | 19 | < .001 |
| Cheek Front - Eyebrow Front | 0.04 | [-0.38, 0.47] | 0.33 | 19 | .999 |
| Cheek Front - Face Side | 0.09 | [-0.14, 0.31] | 1.19 | 19 | .838 |
| Cheek Front - Cheek Side | 0.75 | [0.35, 1.15] | 5.93 | 19 | < .001 |

| Contrast | $\Delta M$ | 95% $\mathrm{CI}_{\mathrm{Tukey}(6)}$ | $t$ | $df$ | $p_{\mathrm{Tukey}(6)}$ |
|---|---|---|---|---|---|
| Cheek Front - Eyebrow Side | 0.70 | [0.22, 1.19] | 4.58 | 19 | .002 |
| Eyebrow Front - Face Side | 0.04 | [-0.39, 0.47] | 0.30 | 19 | > .999 |
| Eyebrow Front - Cheek Side | 0.71 | [0.19, 1.22] | 4.30 | 19 | .004 |
| Eyebrow Front - Eyebrow Side | 0.66 | [0.44, 0.88] | 9.43 | 19 | < .001 |
| Face Side - Cheek Side | 0.66 | [0.37, 0.96] | 7.06 | 19 | < .001 |
| Face Side - Eyebrow Side | 0.62 | [0.15, 1.08] | 4.21 | 19 | .005 |
| Cheek Side - Eyebrow Side | -0.05 | [-0.57, 0.48] | -0.27 | 19 | > .999 |

**Exp. 1: Front-Side Glossiness**

Table S5. *Summary of two-way repeated-measures ANOVA of Rating in Front-Side (Glossiness)*

| Effect | $\hat{\eta}_p^2$ | $F$ | $df^{\mathrm{GG}}$ | $df_{\mathrm{res}}^{\mathrm{GG}}$ | $MSE$ | $p$ |
|---|---|---|---|---|---|---|
| Part | .219 | 5.33 | 1.55 | 29.47 | 0.55 | .016 |
| Direction | .852 | 109.65 | 1 | 19 | 0.10 | < .001 |
| Part × Direction | .503 | 19.25 | 1.52 | 28.79 | 0.09 | < .001 |

Table S6. *Post hoc comparisons - Part*

| Contrast | $\Delta M$ | 95% $\mathrm{CI}_{\mathrm{Tukey}(3)}$ | $t$ | $df$ | $p_{\mathrm{Tukey}(3)}$ |
|---|---|---|---|---|---|
| Face - Cheek | 0.25 | [-0.01, 0.52]$^{a}$ | 2.44 | 19 | .060$^{a}$ |

| Contrast | $\Delta M$ | 95% $\mathrm{CI}_{\mathrm{Tukey}(3)}$ | $t$ | $df$ | $p_{\mathrm{Tukey}(3)}$ |
|---|---|---|---|---|---|
| Face - Eyebrow | 0.48 | [0.09, 0.86]$^b$ | 3.15 | 19 | .014$^b$ |
| Cheek - Eyebrow | 0.22 | [-0.22, 0.67]$^c$ | 1.28 | 19 | .423$^c$ |

Table S7. *Post hoc comparisons - Direction*

| Contrast | $\Delta M$ | 95% CI | $t$ | $df$ | $p$ |
|---|---|---|---|---|---|
| Front - Side | 0.61 | [0.49, 0.73] | 10.47 | 19 | < .001 |

Table S8. *Post hoc comparisons - Part * Direction*

| Contrast | $\Delta M$ | 95% $\mathrm{CI}_{\mathrm{Tukey}(6)}$ | $t$ | $df$ | $p_{\mathrm{Tukey}(6)}$ |
|---|---|---|---|---|---|
| Face Front - Cheek Front | -0.09 | [-0.50, 0.33] | -0.67 | 19 | .983 |
| Face Front - Eyebrow Front | 0.41 | [-0.06, 0.87] | 2.78 | 19 | .104 |
| Face Front - Face Side | 0.34 | [0.22, 0.45] | 9.25 | 19 | < .001 |
| Face Front - Cheek Side | 0.93 | [0.55, 1.32] | 7.67 | 19 | < .001 |
| Face Front - Eyebrow Side | 0.88 | [0.37, 1.40] | 5.41 | 19 | < .001 |
| Cheek Front - Eyebrow Front | 0.50 | [-0.07, 1.07] | 2.76 | 19 | .109 |
| Cheek Front - Face Side | 0.42 | [-0.01, 0.86] | 3.12 | 19 | .054 |
| Cheek Front - Cheek Side | 1.02 | [0.58, 1.46] | 7.37 | 19 | < .001 |
| Cheek Front - Eyebrow Side | 0.97 | [0.30, 1.65] | 4.54 | 19 | .003 |

| Contrast | $\Delta M$ | 95% $\mathrm{CI}_{\mathrm{Tukey}(6)}$ | $t$ | $df$ | $p_{\mathrm{Tukey}(6)}$ |
|---|---|---|---|---|---|
| Eyebrow Front - Face Side | -0.07 | [-0.53, 0.39] | -0.50 | 19 | .996 |
| Eyebrow Front - Cheek Side | 0.52 | [-0.02, 1.06] | 3.07 | 19 | .060 |
| Eyebrow Front - Eyebrow Side | 0.48 | [0.29, 0.66] | 8.25 | 19 | < .001 |
| Face Side - Cheek Side | 0.60 | [0.23, 0.96] | 5.21 | 19 | < .001 |
| Face Side - Eyebrow Side | 0.55 | [0.02, 1.07] | 3.30 | 19 | .037 |
| Cheek Side - Eyebrow Side | -0.05 | [-0.66, 0.56] | -0.25 | 19 | > .999 |

**Exp. 1: Front-Side Attractiveness**

Table S9. *Summary of two-way repeated-measures ANOVA of Rating in Front-Side (Attractiveness)*

| Effect | $\hat{\eta}^2_p$ | $F$ | $df^{\mathrm{GG}}$ | $df^{\mathrm{GG}}_{\mathrm{res}}$ | $MSE$ | $p$ |
|---|---|---|---|---|---|---|
| Part | .264 | 6.81 | 2.00 | 37.94 | 0.27 | .003 |
| Direction | .893 | 157.78 | 1 | 19 | 0.04 | < .001 |
| Part × Direction | .551 | 23.34 | 1.36 | 25.91 | 0.08 | < .001 |

Table S10. *Post hoc comparisons - Part*

| Contrast | $\Delta M$ | 95% $\mathrm{CI}_{\mathrm{Tukey}(3)}$ | $t$ | $df$ | $p_{\mathrm{Tukey}(3)}$ |
|---|---|---|---|---|---|
| Face - Cheek | 0.15 | [-0.15, 0.45]$^a$ | 1.27 | 19 | .427$^a$ |

| Contrast | $\Delta M$ | 95% $CI_{Tukey(3)}$ | $t$ | $df$ | $p_{Tukey(3)}$ |
|---|---|---|---|---|---|
| Face - Eyebrow | 0.42 | [0.12, 0.72]$^{b}$ | 3.60 | 19 | .005$^{b}$ |
| Cheek - Eyebrow | 0.27 | [-0.02, 0.56]$^{c}$ | 2.40 | 19 | .066$^{c}$ |

Table S11. *Post hoc comparisons - Direction*

| Contrast | $\Delta M$ | 95% CI | $t$ | $df$ | $p$ |
|---|---|---|---|---|---|
| Front - Side | 0.47 | [0.39, 0.55] | 12.56 | 19 | < .001 |

Table S12. *Post hoc comparisons - Part * Direction*

| Contrast | $\Delta M$ | 95% $CI_{Tukey(6)}$ | $t$ | $df$ | $p_{Tukey(6)}$ |
|---|---|---|---|---|---|
| Face Front - Cheek Front | 0.02 | [-0.31, 0.36] | 0.21 | 19 | > .999 |
| Face Front - Eyebrow Front | 0.06 | [-0.31, 0.44] | 0.54 | 19 | .994 |
| Face Front - Face Side | 0.15 | [0.02, 0.28] | 3.52 | 19 | .024 |
| Face Front - Cheek Side | 0.42 | [-0.03, 0.87] | 2.98 | 19 | .072 |
| Face Front - Eyebrow Side | 0.93 | [0.53, 1.32] | 7.45 | 19 | < .001 |
| Cheek Front - Eyebrow Front | 0.04 | [-0.32, 0.40] | 0.36 | 19 | .999 |
| Cheek Front - Face Side | 0.12 | [-0.23, 0.48] | 1.10 | 19 | .874 |
| Cheek Front - Cheek Side | 0.40 | [0.12, 0.67] | 4.57 | 19 | .002 |
| Cheek Front - Eyebrow Side | 0.90 | [0.50, 1.30] | 7.16 | 19 | < .001 |

| Contrast | $\Delta M$ | 95% $\mathrm{CI}_{\mathrm{Tukey}(6)}$ | $t$ | $df$ | $p_{\mathrm{Tukey}(6)}$ |
|---|---|---|---|---|---|
| Eyebrow Front - Face Side | 0.08 | [-0.32, 0.49] | 0.64 | 19 | .986 |
| Eyebrow Front - Cheek Side | 0.36 | [-0.02, 0.73] | 3.01 | 19 | .068 |
| Eyebrow Front - Eyebrow Side | 0.86 | [0.62, 1.11] | 11.03 | 19 | < .001 |
| Face Side - Cheek Side | 0.27 | [-0.17, 0.72] | 1.95 | 19 | .405 |
| Face Side - Eyebrow Side | 0.78 | [0.37, 1.19] | 6.02 | 19 | < .001 |
| Cheek Side - Eyebrow Side | 0.50 | [0.03, 0.97] | 3.39 | 19 | .031 |

**Exp. 1: Nomake-Make Moisture**

Table S13. *Summary of two-way repeated-measures ANOVA of Rating in Nomake-Make (Moisuture)*

| Effect | $\hat{\eta}_p^2$ | $F$ | $df^{\mathrm{GG}}$ | $df_{\mathrm{res}}^{\mathrm{GG}}$ | $MSE$ | $p$ |
|---|---|---|---|---|---|---|
| Part | .336 | 9.61 | 1.28 | 24.33 | 0.42 | .003 |
| Make | .143 | 3.18 | 1 | 19 | 0.08 | .091 |
| Part × Make | .533 | 21.70 | 1.35 | 25.73 | 0.05 | < .001 |

Table S14. *Post hoc comparisons - Part*

| Contrast | $\Delta M$ | 95% $\mathrm{CI}_{\mathrm{Tukey}(3)}$ | $t$ | $df$ | $p_{\mathrm{Tukey}(3)}$ |
|---|---|---|---|---|---|
| Face - Cheek | 0.44 | [0.29, 0.59]$^{a}$ | 7.56 | 19 | < .001$^{a}$ |

| Contrast | $\Delta M$ | 95% $CI_{Tukey(3)}$ | $t$ | $df$ | $p_{Tukey(3)}$ |
|---|---|---|---|---|---|
| Face - Eyebrow | 0.44 | [0.10, 0.77]$^b$ | 3.30 | 19 | .010$^b$ |
| Cheek - Eyebrow | 0.00 | [-0.35, 0.35]$^c$ | 0.00 | 19 | > .999$^c$ |

Table S15. *Post hoc comparisons - Part * Make*

| Contrast | $\Delta M$ | 95% $CI_{Tukey(6)}$ | $t$ | $df$ | $p_{Tukey(6)}$ |
|---|---|---|---|---|---|
| Face Nomake - Cheek Nomake | 0.38 | [0.19, 0.56] | 6.37 | 19 | < .001 |
| Face Nomake - Eyebrow Nomake | 0.17 | [-0.22, 0.56] | 1.36 | 19 | .747 |
| Face Nomake - Face Make | -0.31 | [-0.47, -0.16] | -6.50 | 19 | < .001 |
| Face Nomake - Cheek Make | 0.19 | [-0.05, 0.43] | 2.45 | 19 | .188 |
| Face Nomake - Eyebrow Make | 0.39 | [-0.04, 0.82] | 2.87 | 19 | .088 |
| Cheek Nomake - Eyebrow Nomake | -0.21 | [-0.64, 0.23] | -1.51 | 19 | .662 |
| Cheek Nomake - Face Make | -0.69 | [-0.89, -0.48] | -10.66 | 19 | < .001 |
| Cheek Nomake - Cheek Make | -0.19 | [-0.34, -0.04] | -4.07 | 19 | .007 |
| Cheek Nomake - Eyebrow Make | 0.02 | [-0.41, 0.44] | 0.12 | 19 | > .999 |
| Eyebrow Nomake - Face Make | -0.48 | [-0.98, 0.01] | -3.09 | 19 | .057 |
| Eyebrow Nomake - Cheek Make | 0.02 | [-0.52, 0.55] | 0.10 | 19 | > .999 |
| Eyebrow Nomake - Eyebrow Make | 0.22 | [-0.11, 0.55] | 2.13 | 19 | .314 |
| Face Make - Cheek Make | 0.50 | [0.29, 0.71] | 7.50 | 19 | < .001 |
| Face Make - Eyebrow Make | 0.71 | [0.20, 1.21] | 4.45 | 19 | .003 |

| Contrast | $\Delta M$ | 95% $\mathrm{CI}_{\mathrm{Tukey}(6)}$ | $t$ | $df$ | $p_{\mathrm{Tukey}(6)}$ |
|---|---|---|---|---|---|
| Cheek Make - Eyebrow Make | 0.21 | [-0.28, 0.69] | 1.34 | 19 | .760 |

**Exp. 1: Nomake-Make Glossiness**

Table S16. *Summary of two-way repeated-measures ANOVA of Rating in Nomake-Make (Glossiness)*

| Effect | $\hat{\eta}_p^2$ | $F$ | $df^{\mathrm{GG}}$ | $df_{\mathrm{res}}^{\mathrm{GG}}$ | $MSE$ | $p$ |
|---|---|---|---|---|---|---|
| Part | .219 | 5.33 | 1.55 | 29.47 | 0.55 | .016 |
| Make | .152 | 3.40 | 1 | 19 | 0.08 | .081 |
| Part × Make | .517 | 20.35 | 1.50 | 28.59 | 0.07 | < .001 |

Table S17. *Post hoc comparisons - Part*

| Contrast | $\Delta M$ | 95% $\mathrm{CI}_{\mathrm{Tukey}(3)}$ | $t$ | $df$ | $p_{\mathrm{Tukey}(3)}$ |
|---|---|---|---|---|---|
| Face - Cheek | 0.25 | [-0.01, 0.52]$^{a}$ | 2.44 | 19 | .060$^{a}$ |
| Face - Eyebrow | 0.48 | [0.09, 0.86]$^{b}$ | 3.15 | 19 | .014$^{b}$ |
| Cheek - Eyebrow | 0.22 | [-0.22, 0.67]$^{c}$ | 1.28 | 19 | .423$^{c}$ |

Table S18. *Post hoc comparisons - Part * Make*

| Contrast | $\Delta M$ | 95% $\mathrm{CI}_{\mathrm{Tukey}(6)}$ | $t$ | $df$ | $p_{\mathrm{Tukey}(6)}$ |
|---|---|---|---|---|---|
| Face Nomake - Cheek Nomake | 0.29 | [-0.05, 0.63] | 2.69 | 19 | .123 |
| Face Nomake - Eyebrow Nomake | 0.22 | [-0.27, 0.70] | 1.41 | 19 | .720 |
| Face Nomake - Face Make | -0.24 | [-0.46, -0.02] | -3.50 | 19 | .025 |
| Face Nomake - Cheek Make | -0.03 | [-0.38, 0.33] | -0.23 | 19 | > .999 |
| Face Nomake - Eyebrow Make | 0.50 | [0.05, 0.94] | 3.50 | 19 | .025 |
| Cheek Nomake - Eyebrow Nomake | -0.08 | [-0.65, 0.50] | -0.41 | 19 | .998 |
| Cheek Nomake - Face Make | -0.53 | [-0.89, -0.18] | -4.74 | 19 | .002 |
| Cheek Nomake - Cheek Make | -0.32 | [-0.42, -0.21] | -9.58 | 19 | < .001 |
| Cheek Nomake - Eyebrow Make | 0.21 | [-0.35, 0.76] | 1.17 | 19 | .846 |
| Eyebrow Nomake - Face Make | -0.46 | [-1.05, 0.13] | -2.44 | 19 | .191 |
| Eyebrow Nomake - Cheek Make | -0.24 | [-0.85, 0.37] | -1.25 | 19 | .809 |
| Eyebrow Nomake - Eyebrow Make | 0.28 | [-0.07, 0.63] | 2.56 | 19 | .157 |
| Face Make - Cheek Make | 0.22 | [-0.13, 0.57] | 1.95 | 19 | .403 |
| Face Make - Eyebrow Make | 0.74 | [0.19, 1.29] | 4.25 | 19 | .005 |
| Cheek Make - Eyebrow Make | 0.52 | [-0.06, 1.10] | 2.85 | 19 | .092 |

**Exp. 1: Nomake-Make Attractiveness**

Table S19. *Summary of two-way repeated-measures ANOVA of Rating in Nomake-Make (Attractiveness)*

| Effect | $\hat{\eta}_p^2$ | $F$ | $df^{\text{GG}}$ | $df_{\text{res}}^{\text{GG}}$ | $MSE$ | $p$ |
|---|---|---|---|---|---|---|
| Part | .264 | 6.81 | 2.00 | 37.94 | 0.27 | .003 |
| Make | .831 | 93.21 | 1 | 19 | 0.04 | < .001 |
| Part × Make | .079 | 1.62 | 1.44 | 27.42 | 0.04 | .217 |

Table S20. *Post hoc comparisons - Part*

| Contrast | $\Delta M$ | 95% $\text{CI}_{\text{Tukey(3)}}$ | $t$ | $df$ | $p_{\text{Tukey(3)}}$ |
|---|---|---|---|---|---|
| Face - Cheek | 0.15 | [-0.15, 0.45]$^{a}$ | 1.27 | 19 | .427$^{a}$ |
| Face - Eyebrow | 0.42 | [0.12, 0.72]$^{b}$ | 3.60 | 19 | .005$^{b}$ |
| Cheek - Eyebrow | 0.27 | [-0.02, 0.56]$^{c}$ | 2.40 | 19 | .066$^{c}$ |

Table S21. *Post hoc comparisons - Make*

| Contrast | $\Delta M$ | 95% CI | $t$ | $df$ | $p$ |
|---|---|---|---|---|---|
| Nomake - Make | -0.35 | [-0.42, -0.27] | -9.65 | 19 | < .001 |

Table S22. *Physical quantities and measuring equipment*

| Physical quantity | Item name | Measuring equipment | Unit |
|---|---|---|---|
| Trans epidermal water loss | TEWL | VAPO SCAN AS-VT100RS (Asahi Techno Lab) | g/h・$m^2$ |
| Stratum corneum water content | SKICON | SKICON-200EX-USB (ISB) | $\mu$ S |
| Stratum corneum water content | CORNEO | Corneometer CM825 (Courage+Khazaka electronic) | Arbitrary Unit |
| Oiliness | OIL | Sebumeter SM815 (Courage+Khazaka electronic) | $\mu$ g/$cm^2$ |

Table S23. *Measuring schedule*

| Day 1 | Day 2 | Day 3 | Day 4 |
|---|---|---|---|
| Base makeup A | | | |
| Photographing | | | |
| Face-washing → 15 minutes acclimatization | | | |
| Photographing → Skin measuring | | | |
| Lotion | Lotion + Serum | Lotion + Serum + Cream | Vaseline |
| Photographing | | | |
| Face-washing → Skin care | | | |
| Base makeup A | Base makeup B | Base makeup C | Base makeup D |
| Photographing | | | |

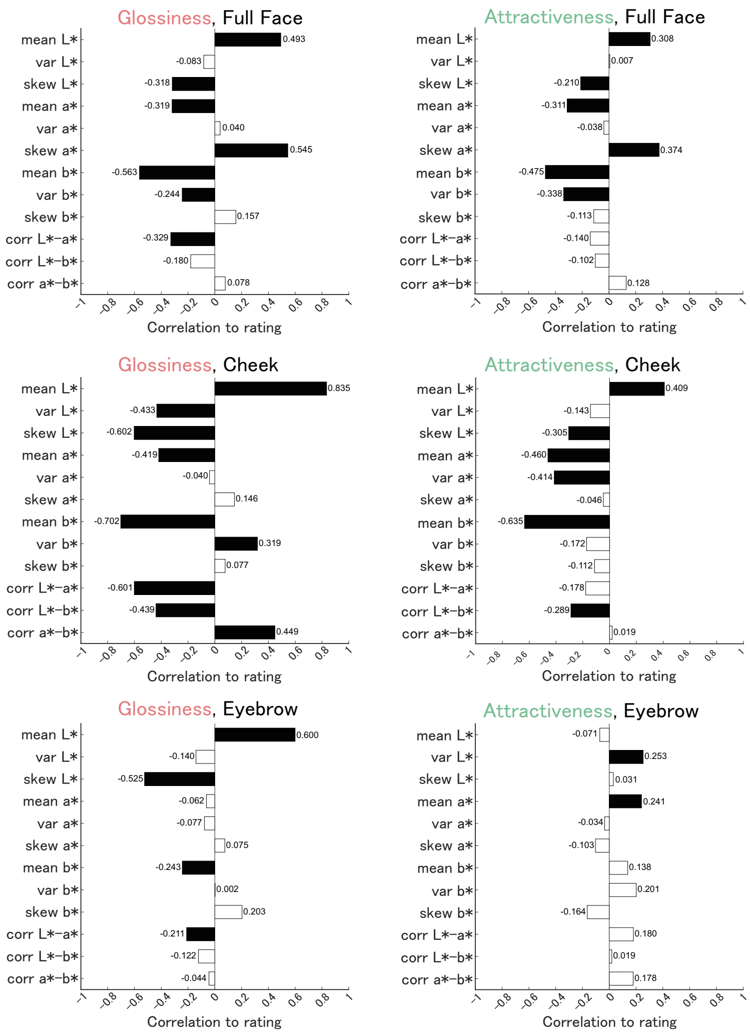


Fig. S1. Correlations between glossiness (left column) and attractiveness (right column) ratings and color statistics in each region (whole face, cheek, eyebrow), respectively.

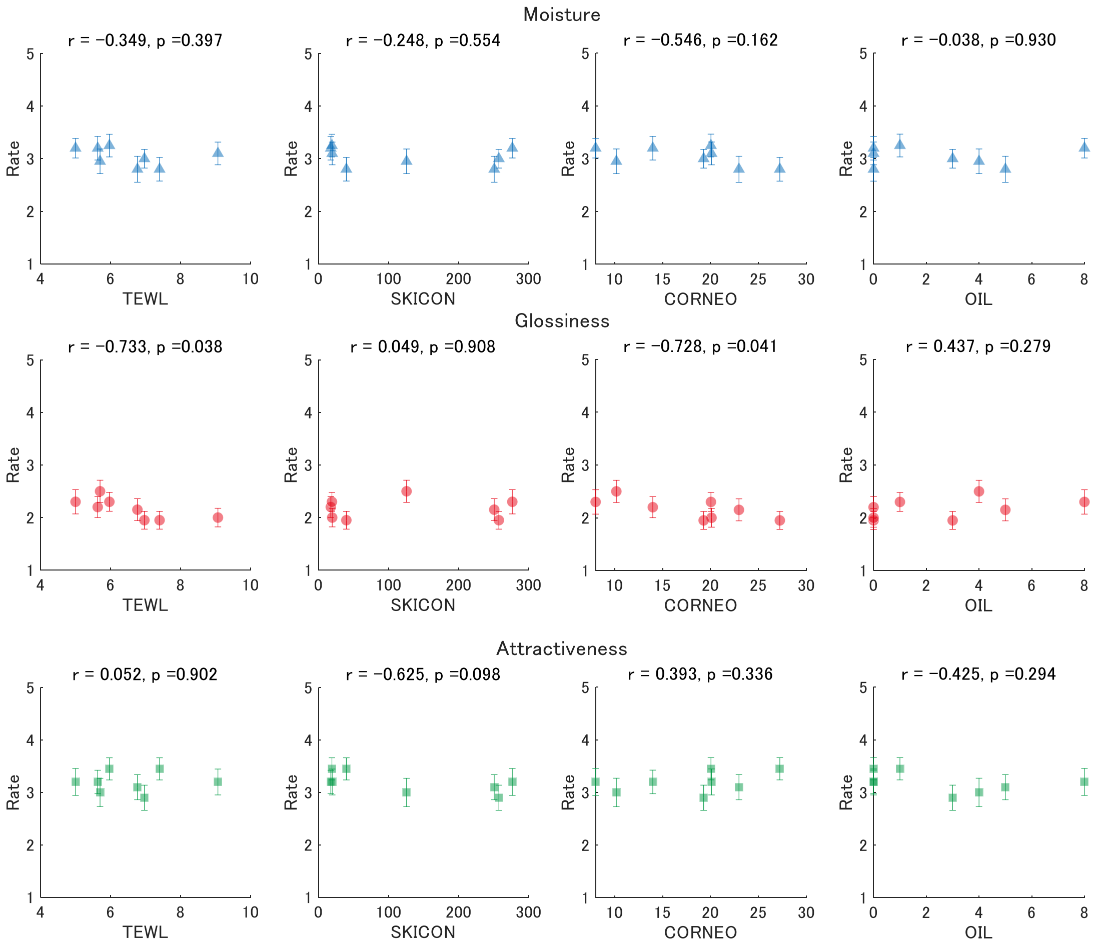


Fig. S2. Scatter plots between physical quantities and ratings.
